# Frontoparietal action-oriented codes support novel instruction implementation

**DOI:** 10.1101/830067

**Authors:** Carlos González-García, Silvia Formica, David Wisniewski, Marcel Brass

**Affiliations:** Department of Experimental Psychology, Ghent University, Belgium

**Keywords:** “Cognitive Control”, “Instructions”, “fMRI”, “multivariate analysis”, “frontoparietal network”, “retro-cues”

## Abstract

A key aspect of human cognitive flexibility concerns the ability to convert complex symbolic instructions into novel behaviors. Previous research proposes that this transformation is supported by two neurocognitive states: an initial declarative maintenance of task knowledge, and an implementation state necessary for optimal task execution. Furthermore, current models predict a crucial role of frontal and parietal brain regions in this process. However, whether declarative and procedural signals independently contribute to implementation remains unknown. We report the results of an fMRI experiment in which participants executed novel instructed stimulus-response associations. We then used a pattern-tracking procedure to quantify the contribution of format-unique signals during instruction implementation. This revealed independent procedural and declarative representations of novel S-Rs in frontoparietal areas, prior to execution. Critically, the degree of procedural activation predicted subsequent behavioral performance. Altogether, our results suggest an important contribution of frontoparietal regions to the neural architecture that regulates cognitive flexibility.

## INTRODUCTION

Instruction following constitutes a powerful instance of human cognitive flexibility. The greater specificity and efficiency in the transmission of task procedures compared to trial-and-error or reinforcement learning make it a distinctive skill that separates humans from other species (Cole et al., 2013). While recent years have witnessed substantial progress in our understanding of instruction following, the precise neural coding schemes that organize brain activity during the rapid transformation of abstract instructed content into effective behavior are still poorly understood.

Previous behavioral studies have reported an intriguing signature of instruction processing, namely, a rapid configuration of instructed content predominantly towards action (González-García et al., 2020; Liefooghe et al., 2012, 2013; Liefooghe and De Houwer, 2018; Meiran et al., 2012, 2015a). This signature separates instruction implementation from related work in task switching and working memory: although preparation for action is not unique to novel instructions, in other contexts repetitive task execution makes it possible to retrieve specific long-term memory traces that allows for successful execution (Qiao et al., 2017; Zhang et al., 2013). In instruction implementation, however, long-term memory traces are reasonably ruled out (Liefooghe et al., 2012; Meiran et al., 2015a; Muhle-Karbe et al., 2016), and rather, an efficient proactive configuration can be achieved without prior practice. This configuration has a profound impact on brain activity. The intention to execute an instruction induces automatic motor activation (Everaert et al., 2014; Meiran et al., 2015b) and specific oscillatory features (Formica et al., 2020b), engages different brain regions to coordinate novel stimuli and responses (Demanet et al., 2016; González-García et al., 2017a; Hartstra et al., 2011; Palenciano et al., 2019b, 2019a), and alters the neural representation of instructed content in control brain regions, primarily, the frontoparietal network (FPN) (Bourguignon et al., 2018; Muhle-Karbe et al., 2017). These and other findings propose a crucial role of the FPN in the rapid access to and configuration of an implementation stage, a highly efficient task readiness state that support successful execution (Bourguignon et al., 2018; González-García et al., 2017b; Hartstra et al., 2011; Muhle-Karbe et al., 2017; Palenciano et al., 2019b, 2019a; Woolgar et al., 2015).

To account for these findings, prominent theoretical models (Brass et al., 2017) put forward a *serial-coding hypothesis* of frontoparietal function, a multi-step process in which the FPN first encodes instructed information into a *declarative* code, that is, a persistent representation of the memoranda conveyed by the instruction. When this information becomes behaviorally relevant, FPN declarative representations are transformed into an implementation state that is optimized for specific task demands (Brass et al., 2017). Current models propose that such implementation state consists primarily of *procedural* codes, a proactive binding of relevant perceptual and motor information into a compound representation that leads to the boost of relevant action codes related to behavioral routines (Muhle-Karbe et al., 2017).

However, the characterization of neural coding during implementation remains unclear, primarily due to the fact that previous analytical approaches were unable to track representational formats of specific nature. Previous work thus identified some properties of the FPN during the implementation of novel instructions, such as enhanced decoding of stimulus (González-García et al., 2017a; Muhle-Karbe et al., 2017) and rule identity (Ruge et al., 2019), or altered similarity within to-be-implemented S-R associations (Bourguignon et al., 2018; Palenciano et al., 2019b). Although these results reveal unique signatures of instruction implementation, they are agnostic regarding the functional representational state underlying such effects, that is, the extent to which they capture a contribution of procedural and declarative signals. Furthermore, previous approaches were not able to eliminate the contribution of domain-general processes, such as arousal or attention, which could potentially drive some of the differences between implementation and other experimental conditions. Therefore, currently, it cannot be discerned whether such implementation state is uniquely supported through the proposed procedural codes, or whether it additionally preserves task content in an independent, declarative format. Furthermore, the specific contribution of these two representational formats to successful behavior remains unknown.

Here, we used a canonical template tracking procedure to capture whether different signals governed FPN activity during the prioritization of novel instructions (Brass et al., 2017). Using data from two independent localizers that encouraged either a declarative or action-oriented maintenance of novel instructions, we derived instruction-specific multivariate patterns of activity in declarative and procedural formats, respectively. We then assessed the contribution of these canonical declarative and procedural templates prior to task execution. Importantly, this partialling logic allowed to determine the format-specific contribution of procedural and declarative representational formats and to partial out the contribution of domain-general processes.

## MATERIALS AND METHODS

Methods are reported, when applicable, in accordance with the Committee on Best Practices in Data Analysis and Sharing (COBIDAS) report (Nichols et al., 2017).

### Participants

Thirty-two participants (mean age = 23.16, range = 19-33; 20 females, 12 males) recruited from the participants’ pool from Ghent University participated in exchange of 40 euros. They were all right-handed (confirmed by the Edinburgh handedness inventory), clinically healthy and MRI-safe. The study was approved by the UZ Ghent Ethics Committee and all participants provided informed consent before starting the experiment. Of the initial 32 participants, 3 were excluded after acquisition (1 participant performed at chance during the task; 1 participant had an error rate of 1 in catch trials (see below); 1 participant’s within-run head movement exceeded voxel size), resulting in a final sample of 29 participants. Due to an incomplete orthogonalization of the cued and uncued S-R pairings, the first three participants were excluded from multivariate analyses (n = 26).

### Apparatus, stimuli, and procedure

S-R associations were created by combining images with words that indicated the response finger. Each S-R association was presented just once during the entire experiment to prevent the formation of long-term memory traces (Meiran et al., 2015a). Given this prerequisite, images of animate (non-human animals) and inanimate (vehicles and instruments) items were compiled from different available databases (Brady et al., 2013, 2008; Brodeur et al., 2014; Griffin et al., 2006; Konkle et al., 2010), creating a pool of 1550 unique pictures (770 animate items, 780 inanimate). To increase perceptual similarity and facilitate recognition, the background was removed from all images, items were centered in the canvas, and images were converted to black and white.

The response dimension was defined by the combination of a word (“index” or “middle”) and the position of the mapping in the encoding screen. For instance, if an S-R pair containing the word “index” was displayed on the left-hand side of the screen, this informed participants that the correct response associated with that particular stimulus would be “*left* index”. This allowed us to have 2 mappings on screen that involved the same (stimulus and) *response category* (e.g. index finger) but different effectors (e.g. *left* index finger vs *right* index finger).

Importantly, even though specific S-R associations were presented only once throughout the experiment, they could be grouped depending on the specific combination of stimulus and response dimensions. Specifically, the combination of the 2 stimulus dimensions (animate/inanimate items) and the 2 response dimensions (index/middle finger) lead to 4 finger-animacy pairings: S-R 1 (animate-index), S-R 2 (inanimate-index), S-R 3 (animate-middle), and S-R 4 (inanimate-middle).

In the main task, each trial started with an encoding screen (5000 ms) that displayed 4 S-R associations. The two mappings on the upper half of the encoding screen belonged to one S-R pairing, and the other two belonged to another S-R pairing. Immediately after the encoding screen, a retro-cue appeared. Informative retro-cues (75% of trials) consisted of an arrow centered in the middle of the screen pointing either upwards or downwards. Therefore, informative retro-cues did not select a specific S-R mapping but rather two mappings belonging to the same S-R pairing (e.g. “animate - index finger”). Such grouping was crucial for analysis purposes since it allowed us to identify the *selected* S-R pairing, as well as the *unselected* category that was initially encoded but could be dropped from working memory after the retro-cue. Additionally, for each trial, we identified the *not presented* S-R pairings, which would serve as empirical baseline for our template tracking analysis (see below). In contrast, neutral retro-cues did not select any mapping. The retro-cue was displayed for 1000 ms and was followed by a fixation point (cue-target interval; CTI), which duration was jittered following a pseudo-logarithmic distribution (mean duration = 2266 ms, SD = 1276 ms, range = [600-5000]). Directly after the CTI, a target was on screen for 1500 ms. Target screens displayed the image belonging to one of the selected mappings, prompting participants to execute the associated response by pressing the corresponding button in an MRI-compatible button box. In neutral trials, the target could be the stimulus of any of the 4 S-R encoded mappings. Additionally, in ~6% of trials, a catch target appeared. This consisted of a new image, different from any of the encoded stimuli, to which participants had to answer by pressing the 4 available buttons in the response box. Catch trials were included to ensure that participant encoded all four S-R associations and were equally likely after an informative and a neutral retro-cue. Last, after the target screen, a fixation point was shown between trials (inter-trial interval, ITI) for a jittered duration (following the same parameters as the CTI jitter). Each trial lasted on average 12 seconds. The sequence of trial events is depicted in Figure 1.

**Figure 1.**
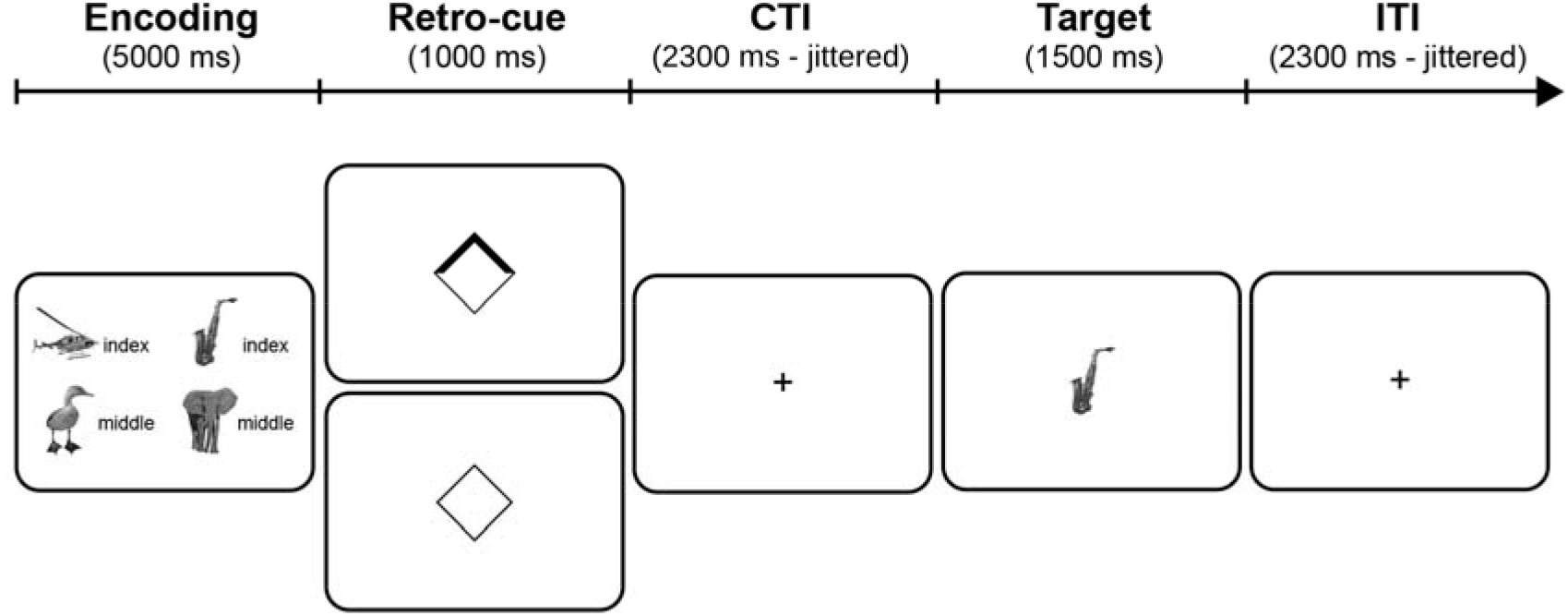
Behavioral paradigm. On each trial, participants first encoded four novel S-R mappings consisting in the association between an (animate or inanimate) item and a response (index or middle fingers; response hand defined by the position of the mapping on the screen; e.g. “helicopter-index” on the left-hand side of the screen requested participants to press the *left* index if the target screen displayed a helicopter). After the encoding screen, an informative retro-cue (75% of the trials) signaled the relevance of two of the mappings. In the remaining 25% of trials, a neutral retro-cue appeared, and none of the mappings were cued. Last, a target stimulus prompted participants to provide the associated response (in this example, “right index” finger press).

The main task was divided into 4 runs. Each run contained 51 trials (48 regular and 3 catch trials). Of the 48 regular trials, 75% contained an informative retro-cue, and the remaining trials displayed neutral retro-cues. The S-R pairings selected and unselected by the retro-cue were fully counterbalanced, resulting in 36 trials per pairing across the entire experiment. For instance, there were 36 trials in which Pairing 1 mappings were selected by the retro-cue. Of these 36 trials, in one third, the unselected mappings (that is, mappings shown in the encoding screen but not selected by the retro-cue) belonged to Pairing 2, another third to Pairing 3, and the last third to Pairing 4. Each run lasted around 10 minutes, and the main task, containing 204 trials, lasted around 40 minutes in total. Prior to the main task, outside of the scanner, participants performed a practice session with trials following the same structure described above with the exception that feedback was included to help familiarization. The practice session was structured in blocks of 11 trials. Participants performed these blocks until they achieved at least 9 correct responses. S-R mappings used during the practice were never used again.

After the main task, participants performed two localizers to obtain an independent canonical representation of each S-R pairing in the two formats of interest. The two localizers were aimed at encouraging either a primarily procedural or a primarily declarative coding of new S-Rs. Although a localizer eliciting uniquely one of these two types of coding is hard to conceive (for instance, one could claim a declarative representation of the elements of a task is required before any procedural representation can emerge (Formica et al., 2020a)), our pattern analysis (see below) capitalized on the specific engagement of procedural and declarative strategies encouraged by each of these localizers.

The structure of the task was almost identical in the two localizers. In both localizers, trials started with an encoding screen (2000 ms) that contained two mappings of the same S-R pairing, followed by an inter-stimulus interval of jittered duration (same parameters as the jitters in the main task). Importantly, in both localizers, even though the two encoded mappings belonged to the same S-R pairing, they specified different effectors (for instance: “if you see an elephant, press left index finger; if you see a tiger, press right index finger), and therefore participants needed to maintain both mappings rather using other strategies, such as remembering 2 images and one response.

Last, a target screen appeared (1500 ms) followed by a jittered ITI. The target screen differed in the two localizers and was inspired by previous studies investigating the dissociation of implementing vs. memorizing new instructions (Liefooghe et al., 2012; Liefooghe and De Houwer, 2018; Muhle-Karbe et al., 2017). In the procedural localizer, the target consisted of a single image that prompted participants to execute the associated response. Although this configuration renders the procedural localizer similar to the main task, it remained different in a crucial aspect. Whereas in the procedural localizer participants could prepare for executing one of the 2 mappings directly in the encoding screen, in the main task this highly action-oriented coding format was strategically optimal only after the selection process elicited by the retro-cue. Since our analyses focused on this moment of the main task, the localizer thus provided a means to test whether the selection of a novel S-R from working memory engaged similar procedural signals.

The declarative localizer, in contrast to the procedural one, displayed a memory target consisting of one image and one response finger (e.g. left index). Participants were trained to answer whether the displayed mapping was correct (same association as the encoded one) or incorrect (different association) by pressing both left-hand buttons (when “correct”) or both right-hand buttons (when “incorrect”). Therefore, in the memorization localizer, participants never had to prepare to execute the encoded mapping but rather just maintain its information. As in the main task, catch trials consisted of new images, to which participants had to respond by pressing all 4 available buttons. Each trial lasted around 8 s on average, and each localizer contained 66 trials (15 per S-R pairing + 6 catch trials), resulting in a total of 9 minutes per localizer. Given that the task demands for the procedural localizer were more similar to the main task, this localizer was performed always before the declarative localizer, which required more detailed explanation to participants. Importantly, the nature of our template tracking approach (see below) accounted for any potential order confound in such analysis, since template activation is measured against an empirical within-localizer baseline, and not directly compared between localizers.

All tasks were presented in PsychoPy 2 (Peirce, 2007) running on a Windows PC and back-projected onto a screen located behind the scanner. Participants responded using an MRI-compatible button box on each hand (each button box contained two buttons, on which participants placed their index and middle fingers).

### Data acquisition and preprocessing

Imaging was performed on a 3T Magnetom Trio MRI scanner (Siemens Medical Systems, Erlangen, Germany), equipped with a 64-channel head coil. T1 weighted anatomical images were obtained using a magnetization-prepared rapid acquisition gradient echo (MP-RAGE) sequence (TR=2250 ms, TE=4.18 ms, TI=900 ms, acquisition matrix=256 × 256, FOV=256 mm, flip angle=9°, voxel size=1 × 1 × 1 mm). Moreover, 2 field map images (phase and magnitude) were acquired to correct for magnetic field inhomogeneities (TR=520 ms, TE1=4.92 ms, TE2=7.38 ms, image matrix=70 x 70, FOV=210 mm, flip angle=60°, slice thickness=3 mm, voxel size=3 x 3 x 2.5 mm, distance factor=0%, 50 slices). Whole-brain functional images were obtained using an echo planar imaging (EPI) sequence (TR=1730 ms, TE=30 ms, image matrix=84 × 84, FOV=210 mm, flip angle=66°, slice thickness=2.5 mm, voxel size=2.5 x 2.5 x 2.5 mm, distance factor=0%, 50 slices with slice acceleration factor 2 (Simultaneous Multi-Slice acquisition)). Slices were orientated along the AC-PC line for each subject.

For each run of the main task, 373 volumes were acquired, whereas 330 volumes were acquired during each localizer. In all cases, the first 8 volumes were discarded to allow for (1) signal stabilization, and (2) sufficient learning time for a noise cancellation algorithm (OptoACTIVE, Optoacoustics Ltd, Moshav Mazor, Israel). Before data preprocessing, DICOM images obtained from the scanner were converted into NIfTI files using HeuDiConv (https://github.com/nipy/heudiconv), in order to organize the dataset in accordance with the BIDS format (Gorgolewski et al., 2017). Further data preprocessing was performed in SPM12 (v7487) running on MATLAB R2016b. First, anatomical images were defaced to ensure anonymization. They were later segmented into gray matter, white matter and cerebro-spinal fluid components using SPM default parameters. In this step, we obtained inverse and forward deformation fields to later (1) normalize functional images to the atlas space (forward transformation) and (2) transform ROIs from the atlas on to the individual, native space of each participant (inverse transformation). Regarding functional images, preprocessing included the following steps in the following order: (1) Images were realigned and unwarped to correct for movement artifacts (using the first scan as reference slice) and magnetic field inhomogeneities (using fieldmaps); (2) slice timing correction; (3) coregistration with T1 (intra-subject registration): rigid-body transformation, normalized mutual information cost function; 4^th^ degree B-spline interpolation; (4) registration to MNI space using forward deformation fields from segmentation: MNI 2mm template space, 4^th^ degree B-spline interpolation; and (5) smoothing (8-mm FWHM kernel). Multivariate analyses were conducted on the unsmoothed, individual subject’s functional data space and results were later pooled across participants for region-of-interest analyses.

### Experimental design and statistical analysis

Our main task design consisted of two within-subject factors orthogonally manipulated: retro-cue status (informative vs. neutral) and selected S-R pairing. Regarding behavioral data analyses, we used JASP (JASP Team, 2018) to perform two-tail paired t-tests comparing reaction times and error rates for trials with informative vs. neutral trials (collapsing across selected S-R pairing).

#### General Linear Model (GLM) estimations and mass-univariate analyses

Four GLMs were estimated for each participant in SPM. First, a GLM was used to assess changes in activation magnitude between informative and neutral retro-cues during the main task. A model was constructed including, for each run, regressors for the encoding screen (zero duration), informative/neutral retro-cues (with duration), informative/neutral CTI (with duration), target (zero duration) and ITI interval (with duration). Trials with errors were included as a different regressor that encompassed the total duration of the trial. All regressors were convolved with a hemodynamic response function (HRF). At the population level, parameter estimates of each regressor were entered into a mixed-effects analysis. To correct for multiple comparisons, first we identified individual voxels that passed a ‘height’ threshold of p < 0.001, and then the minimum cluster size was set to the number of voxels corresponding to p < 0.05, FWE-corrected. This combination of thresholds has been shown to control appropriately for false-positives (Eklund et al., 2016). A second GLM was estimated on the non-normalized and unsmoothed main task data for all multivariate analyses. This GLM contained beta estimates that specified the cued/uncued S-R pairings during informative retro-cues. For each participant and run, a model was built including the following regressors: encoding (zero duration), neutral retro-cues (with duration), targets (zero duration), CTI and ITI (with duration). For informative retro-cues, a regressor that encompassed the total duration of the retro-cue was created for each S-R pairing combination (e.g. CuedPairing1_UncuedPairing2), resulting in a total of 12 regressors (3 per finger-animacy pairing). Errors were included as a different regressor encompassing the full duration of the trial. Last, a third and fourth GLMs were performed on the non-normalized and unsmoothed data from the two localizers. For each localizer, we built a model that contained regressors for the encoding screen (zero duration), encoding-target interval (ISI, with duration) for each S-R pairing (total of 4 regressors), target (zero duration), ITI (with duration), and errors (full trial). As in the previous GLM, these models were not used in a population-level GLM and were estimated for later use in the canonical template tracking procedure.

#### Multivariate pattern analysis (MVPA)

MVPA was performed on the beta images of the second GLM using The Decoding Toolbox (Hebart et al., 2015) (v3.99). To assess the representation of cued S-R pairings during implementation, we carried out ROI-based one-vs-one multiclass decoding of S-R pairings. In each fold of the leave-one-run-out procedure, we trained a classifier (linear support vector machine (SVM); regularization parameter = 1) on the identity of the *cued* S-R pairing using all informative retro-cue betas but four (one from each class). The classifier was then tested on the remaining samples. Thus, the held-out data in each cross-validation fold was from different experimental runs to the training data. The accuracy was averaged across folds. Only one decoding was performed per ROI, using all voxels. To remove any potential magnitude difference between classes, we z-scored the values of each condition across voxels before the analysis (therefore, each condition that entered the analysis had a mean activation of 0 and an s.d. of 1). We then repeated the same procedure but now training and testing the classifier on the identity of the *uncued* S-R pairing.

Statistics of decoding analyses followed a permutation approach (Combrisson and Jerbi, 2015). For each ROI, we computed a null distribution by repeating the decoding protocol 1000 times swapping the labels of the true classes. We then established the chance level for a given participant as the mean value of this null distribution. To assess significance at the population level, we first compared accuracy minus chance scores of all participants against 0, using a one-sample t-test. Then, we computed the empirical null distribution of t-values by, on each of 1000 permutations, randomly flipping the sign of each individual score and performing a new t-test. Finally, an effect was considered significant if the observed t-value was larger than 95% of the t-values in the null distribution (thus, significance level = p < 0.05).

#### Canonical template tracking procedure

The main goal of the current study was to assess the contribution of procedural and declarative signals during instruction implementation. To do so, we followed a canonical template tracking procedure (Wimber et al., 2015) (see Figure 4 for a visual representation of the analysis). The main rationale of this analysis was (1) to obtain canonical representations of the different S-R pairings under the two different formats of interest (procedural and declarative) from the ISI of the localizers, and later (2) estimate the extent of variance during implementation in the main task uniquely explained by each of these representations. Importantly, this analysis was aimed at obtaining evidence for the presence (or lack thereof) of procedural and/or declarative signals and not to compare their strengths.

The functional localizers performed after the main task allowed us to obtain a participant-specific canonical pattern of activation for each S-R pairing in declarative and procedural formats. All patterns were derived from beta weights of the GLMs described in the section General Linear Model estimations. Prior to analysis, betas were converted into t-maps and, given the impact of noise on correlation estimates, we performed multivariate noise normalization on each individual run of the main task and template separately (Walther et al., 2016). To do so, we used the residuals of each participant’s GLMs to estimate the noise covariance between voxels. These estimates, regularized by the optimal shrinkage factor (Ledoit and Wolf, 2004), were used to spatially pre-whiten the t-maps.

To measure the contribution of the canonical patterns during the main task, for each region, we computed the semi-partial correlation between the pattern of activity during the retro-cue in the main task and the canonical template of each S-R pairing in the two formats. Semi-partial correlations make it possible to estimate how much *unique* variance the independent variable (e.g. the residuals of regressing the procedural template of one S-R pairing on the declarative template of the same pairing) explains in relation to the total variance in the dependent variable (e.g. activity during the main task), and are thus more practically relevant than partial correlations because they are scaled to the total variability in the dependent variable, rather than to the variance unaccounted for by the rest of variables.

An important feature of the described template tracking approach is that it was optimized to detect whether the two signals of interest were independently accounting for unique variance during implementation, and not to compare the strength of these two signals. Therefore, the raw semi-partial correlation magnitude of each template with the task was of no interest. Only the relative difference between correlation of cued, uncued S-Rs, and the empirical baseline provided by the not-presented S-R was informative for our hypotheses. Since our GLM included different retro-cue regressors depending on the selected S-R pairing, we could obtain a specific activation value for cued, uncued and not-presented pairings. Importantly, semi-partial correlations were used to obtain the amount of variance shared between the main task and a template of an S-R pairing (e.g. in procedural state) that is not explained by the template of that same pairing in the opposite state (e.g. declarative). As such, this approach is sensitive to content-specific signals and rules out the relative contribution of domain general processes that are shared between the two localizers, ensuring that any significant result would only capture the activation of S-R information in a specific format. To statistically test the boost of cued information, we first normalized the semi-partial correlation scores by using Fisher’s z transformation and then performed paired t-tests between the cued, uncued and not-presented S-R pairings activation (FDR-corrected for multiple comparisons).

### Region-of-interest (ROI) definition

Frontoparietal ROIs were obtained from a parcellated map of the multiple-demand network (Fedorenko et al., 2013). Specifically, frontal ROIs comprised the inferior and middle frontal gyrus regions of the map, and parietal ROIs comprised the inferior and superior parietal cortex regions. All ROIs were registered back to the native space of each subject using the inverse deformation fields obtained during segmentation.

We obtained a ventral visual cortex ROI by extracting the following regions in the WFU pickatlas software (http://fmri.wfubmc.edu/software/PickAtlas): bilateral inferior occipital lobe, parahippocampal gyrus, fusiform gyrus, and lingual gyrus (all bilateral and based on AAL definitions). The primary motor cortex ROI was also obtained using WFU pickatlas by extracting the bilateral M1 region.

## RESULTS

### S-R prioritization enhances instruction execution

Analysis of participants’ behavioral performance revealed that retro-cues helped participants in prioritizing novel S-Rs. Specifically, participants were faster (*t*_28,1_ = 13.51, *p* < 0.001, Cohen’s *d* = 2.51; Fig. 2a) and made less errors (*t*_28,1_ = 7.96, *p* < 0.001, Cohen’s *d* = 1.47; Fig. 2b, left panel) in trials with informative retro-cues, compared to neutral. Participants were slower in catch trials compared to informative (*t*_28,1_ = 11.68, *p* < 0.001, Cohen’s *d* = 2.17) and neutral trials (*t*_28,1_ = 3.36, *p* = 0.002, Cohen’s *d* = 0.63). This longer RT probably reflected the requirement to disengage from the encoded S-Rs and respond correctly to the new, non-encoded image. In line with this interpretation, responses to catch images after a neutral retro-cue (M = 981 ms, SD = 122) were slower than after an informative retro-cue (M = 909 ms, SD = 95; *t*_27,1_ = 3.81, *p* < 0.001, Cohen’s *d* = 0.72). This cost of WM load only modulated RTs: error rates for catch trials were lower than in neutral trials (*t*_28,1_ = 4.83, *p* < 0.001, Cohen’s *d* = 0.90), and not significantly different from informative trials (*t* < 1), suggesting that participants were able to detect new images and, therefore, that they successfully encoded all mappings of the encoding screen.

**Figure 2.**
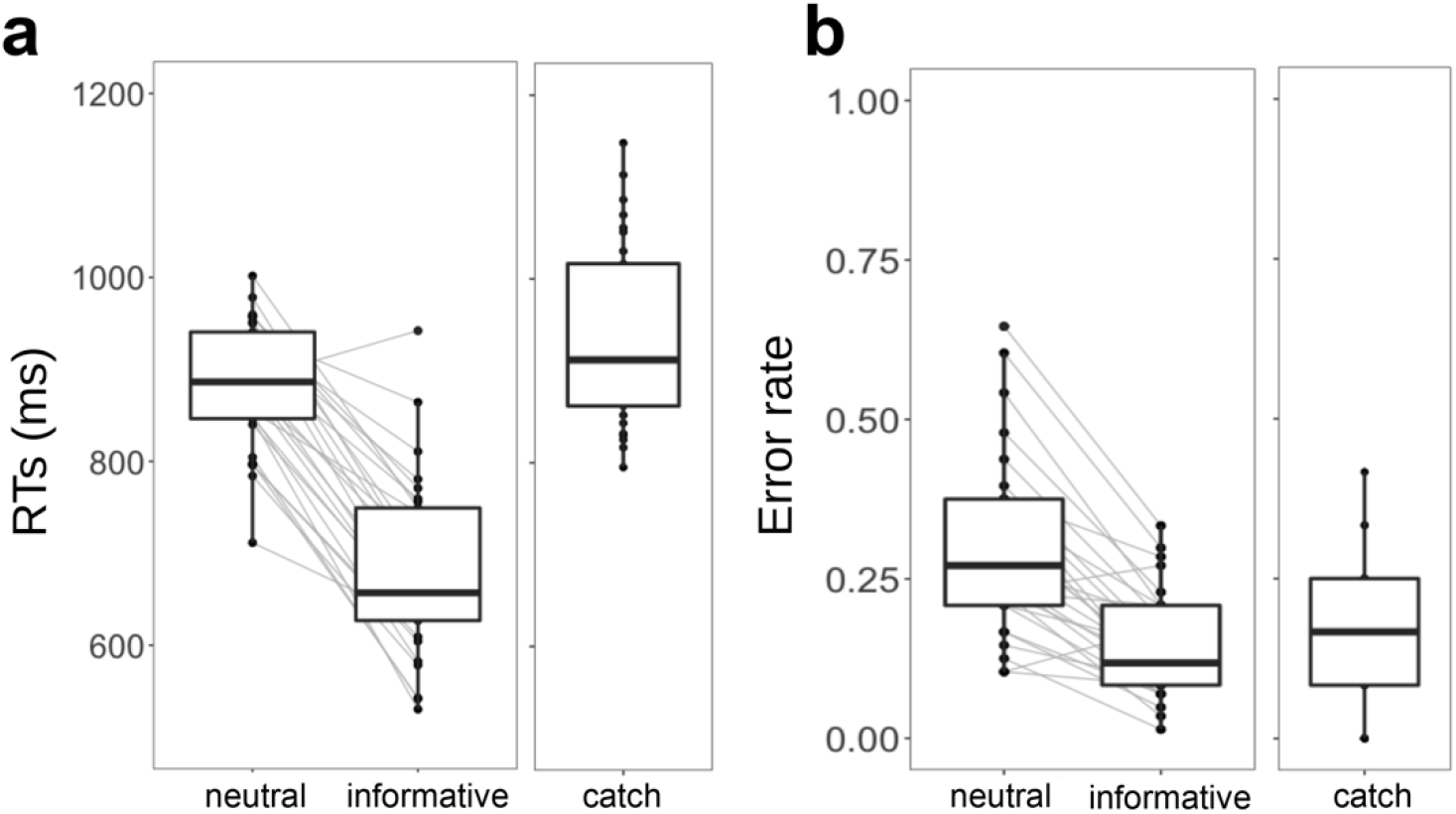
Behavioral results. (**a**) Reaction times in neutral, informative, and catch trials. (**b**) Error rates in neutral, informative, and catch trials. The thick line inside box plots depicts the second quartile (median) of the distribution (n = 29). The bounds of the boxes depict the first and third quartiles of the distribution. Whiskers denote the 1.5 interquartile range of the lower and upper quartile. Dots represent individual subjects’ scores. Grey lines connect dots corresponding to the same participant in two different experimental conditions.

Regarding performance during the two localizers, we expected more successful behavior during the procedural localizer task, given the simpler nature of the task. Accordingly, participants responded faster (t_28,1_ = 25.75, p < 0.001, Cohen’s d = 4.78) and made less errors (t_28,1_ = 3.99, p < 0.001, Cohen’s d = 0.74) during the procedural localizer (RT M = 652 ms, SD = 84; ER M = 0.15, SD = 0.1), compared to declarative one (RT M = 1042, SD = 75; ER M = 0.25, SD = 0.08).

### Identifying novel S-R selection activity

As a first step, we investigated which brain regions were predominantly involved in the selection of instructions from working memory (WM). Based on recent experimental results (González-García et al., 2020; Myers et al., 2018; Yu and Postle, 2018) and theoretical models of WM (Myers et al., 2017), we assumed that selection would prioritize relevant S-R associations into a behavior-optimized state, akin to implementation. As such, retro-cues served as a tool to locate in time the moment after initial encoding in which implementation-specific signals should be magnified in detriment of encoded but uncued S-Rs, which could be potentially dropped from WM. Specifically, we predicted that if such prioritization of S-Rs is indeed similar to instruction implementation, then the FPN should be particularly engaged in trials with informative retro-cues (Bourguignon et al., 2018; González-García et al., 2017a; Jackson and Woolgar, 2018; Muhle-Karbe et al., 2017; Palenciano et al., 2019a; Woolgar et al., 2015). We thus established a set of a priori candidate regions that encompassed frontal (inferior and middle frontal gyri) and (inferior and superior) parietal cortices (see Fig. 3b, and the Region-of-interest definition section in the Methods). We then performed a whole-brain analysis to find regions sensitive to S-R selection (defined as informative vs. neutral retro-cues) in their overall activation magnitude using a general linear model (GLM). We found that informative retro-cues elicited significantly higher activity in regions of the FPN, including the inferior and middle frontal gyri, inferior and superior parietal cortices, as well as regions outside the FPN, such as the lateral occipital cortex (Fig. 3a, primary voxel threshold [*p* < 0.001 uncorrected] and cluster-defining threshold [FWE *p* < .05]). Overall, the resulting statistical map roughly overlap with the set of a priori defined regions of interest (ROIs; Fig. 3b), confirming the involvement of the FPN in S-R selection and, more broadly, providing initial evidence that such prioritization could engage similar mechanisms as instruction implementation.

**Figure 3.**
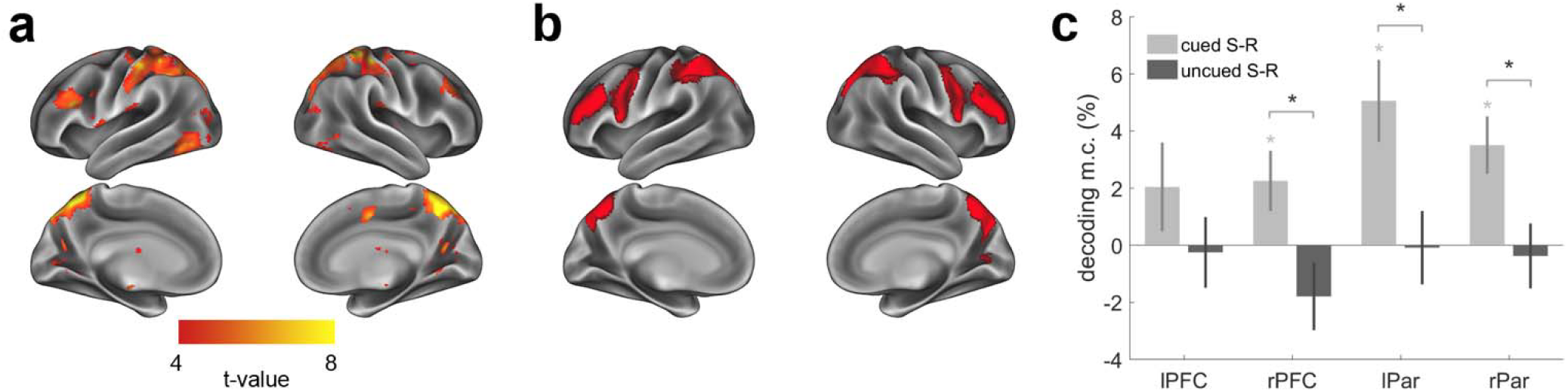
S-R selection induced changes in frontoparietal neural activity. (**a**) GLM contrast of informative > neutral retro-cue trials. Warm colors show regions with significantly higher activity magnitude during informative compared to neutral retro-cues (primary voxel threshold [*p* < 0.001 uncorrected] and cluster-defining threshold [FWE *p* < .05]). (**b**) Set of regions-of-interest defined prior to analyses, encompassing frontal (inferior and middle frontal gyri) and (inferior and superior) parietal cortices. (**c**) Mean S-R pairing decoding (minus empirical chance level) within each region of interest. Error bars denote between-participants s.e.m. Grey asterisks denote accuracies significantly above chance level (permutation-based one-sample t-test, 1k permutations). Black asterisks denote significantly higher accuracies for cued compared to uncued S-R pairings (permutation-based paired t-test, 1k permutations).

Next, we predicted that the prioritization state would modulate the representation of S-R pairings. To test this hypothesis, we performed two similar multivariate decoding analyses in the 4 FPN ROIs. First, we tested if at the moment of the retro-cue the patterns of activity in these four regions carried information about the specific finger-animacy pairing of the cued S-R. We found significant decoding in the right PFC and bilateral parietal ROIs (permutation-based one-sample t-tests, all *p*s < 0.02), and not significant decoding in the left PFC (*t*_25,1_ = 1.69, *p* = 0.1), although a Bayesian t-test suggested no conclusive evidence for neither the alternative nor the null hypothesis in this ROI (BF_10_ = 0.45). Next, we tested the extent to which the FPN also carried information about the encoded, but not cued pairing. In contrast to the previous results, we expected these pairings not to be decodable, given that uncued mappings could be dropped from memory. In line with our prediction, decoding did not reach significance in any of the ROIs (all *p*s > 0.06), and a Bayesian counterpart of the analysis provided support for the null hypothesis (in left DLPFC and bilateral parietal ROIs, all BFs < 0.3) and inconclusive evidence in the right DLPFC (BF_10_ = 0.58). Finally, we directly compared the decoding accuracies for the cued and uncued pairings. This analysis revealed significantly stronger decoding of the cued pairing compared to the uncued one in right PFC and bilateral parietal cortices (permutation-based paired t-tests, all *p*s < 0.02, Fig. 3c; see Table 1 for individual statistics, p-values and BF_10_ estimates).

**Table 1.**
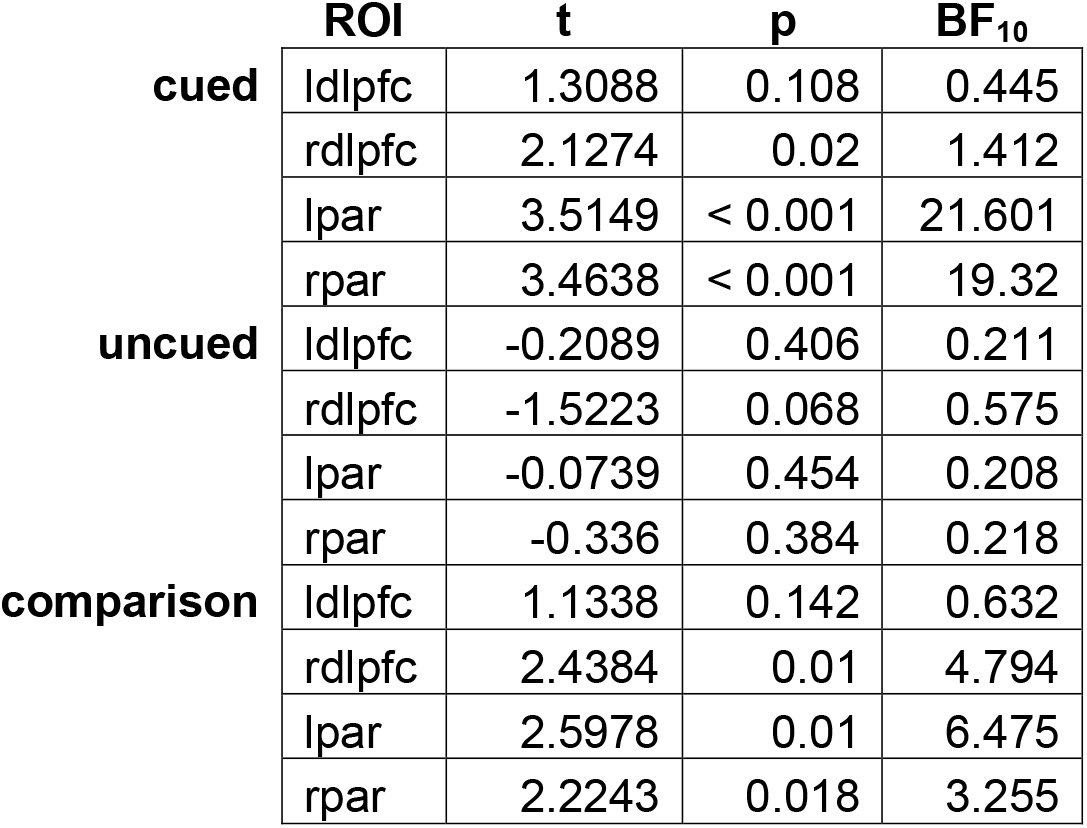
Statistics, p-values and BF_10_ estimates for ROI-based decoding results. BF10 > 3 suggests support for the alternative hypothesis, whereas BF10 < 0.3 indicates support for the null hypothesis.

|  | ROI | t | p | BF<sub>10</sub> |
| --- | --- | --- | --- | --- |
| **cued** | ldlpfc | 1.3088 | 0.108 | 0.445 |
|  | rdlpfc | 2.1274 | 0.02 | 1.412 |
|  | lpar | 3.5149 | < 0.001 | 21.601 |
|  | rpar | 3.4638 | < 0.001 | 19.32 |
| **uncued** | ldlpfc | -0.2089 | 0.406 | 0.211 |
|  | rdlpfc | -1.5223 | 0.068 | 0.575 |
|  | lpar | -0.0739 | 0.454 | 0.208 |
|  | rpar | -0.336 | 0.384 | 0.218 |
| **comparison** | ldlpfc | 1.1338 | 0.142 | 0.632 |
|  | rdlpfc | 2.4384 | 0.01 | 4.794 |
|  | lpar | 2.5978 | 0.01 | 6.475 |
|  | rpar | 2.2243 | 0.018 | 3.255 |

### Tracking format-unique S-R patterns

Altogether, these results show that instruction prioritization has a profound impact on FPN activity, impacting the representation of selected and irrelevant S-Rs. However, similarly to previous studies, they are agnostic regarding the nature of the signals underlying such effect. The main goal of our study was to test whether both declarative and procedural signals contributed to the representational organization of FPN activity during instruction implementation. To do so, we implemented a canonical template tracking procedure that allowed us to estimate neural activations of specific S-R pairings under the two functional formats of interest (see Figure 4, for a visual representation of the procedure, and Methods, for a detailed description of the analysis). Importantly, this approach revealed the amount of shared variance between task data and a given template (e.g. S-R pairing 1 in procedural state) that is not explained by the same template in the alternative state (e.g. S-R pairing 1 in declarative state). Therefore, processes common to both localizers (e.g. arousal, domain-general attention and/or task preparation) cannot inflate correlations, and any significant enhancement from baseline rather reflects the activation of S-R-specific information in a specific format during the main task.

**Figure 4.**
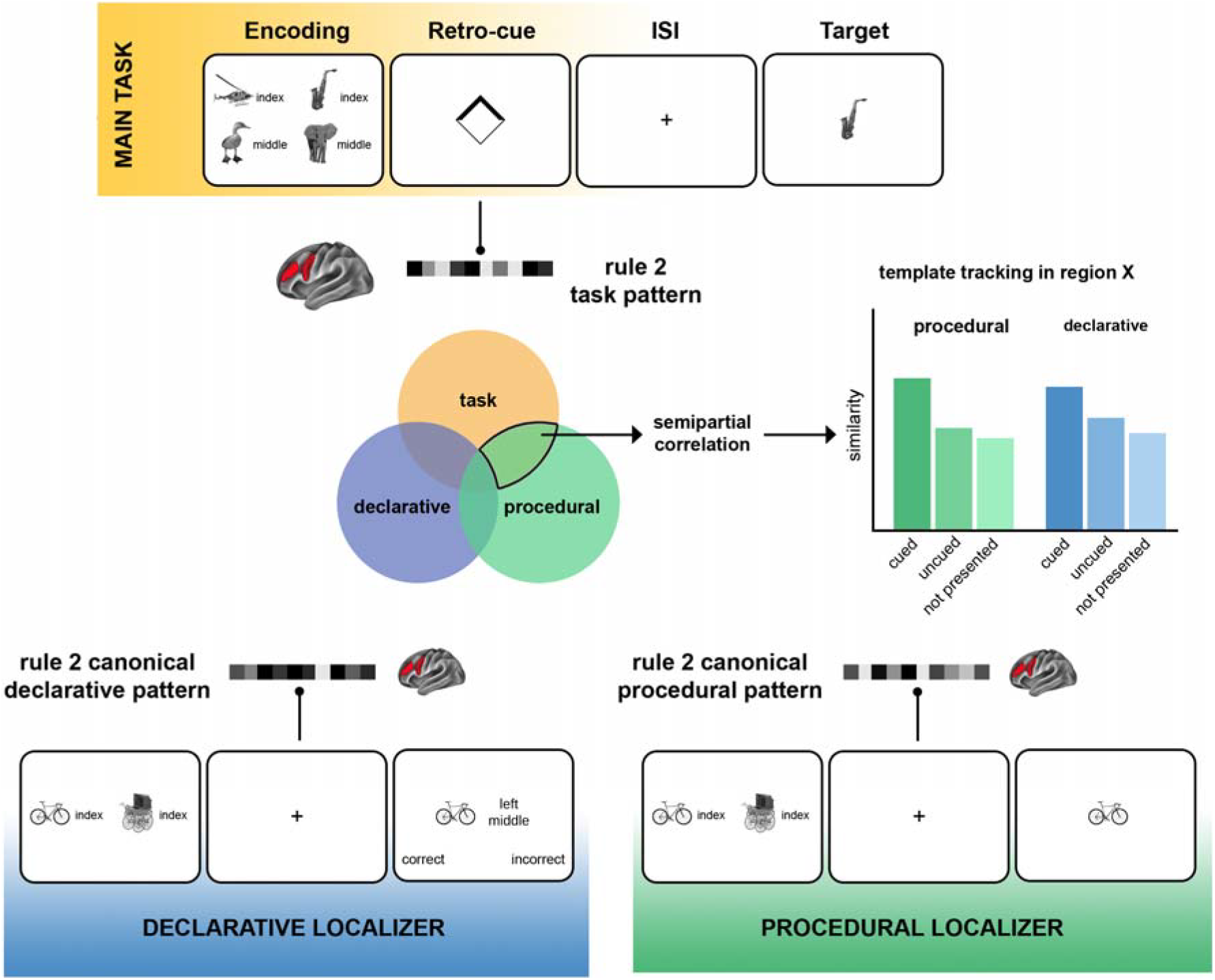
Schematic of the canonical template tracking procedure. For each region of interest, we extracted the pattern of activity of specific S-R pairings during informative retro-cues (upper panel, in yellow) and computed similarity with canonical templates of such pairings in declarative (bottom left, in blue) and procedural (bottom right, in green) formats, obtained in two separate localizers. Importantly, similarity was assessed via semi-partial correlations, obtaining the proportion of uniquely shared variance between task and template data (middle, Venn diagram) of the cued, uncued and not-presented S-R pairings, which provide an empirical baseline. Graphs represent a hypothetical set of results, in which implementation recruits non-overlapping procedural and declarative representations of cued S-R pairing. This informational boost, relative to baseline (not-presented S-R pairings), is superior to that of the uncued pairing.

To validate this procedure outside the FPN, we created an ROI comprising the primary motor cortex, where implementation should be dominated by action-oriented signals and no declarative information about S-R pairings is expected. The results obtained (Fig. 5) matched the predictions, revealing a specific enhancement of procedural information of the cued pairing compared to the uncued (t_25,1_ = 4.08, p < 0.001, Cohen’s d = 0.80), and critically, to the empirical baseline defined by the not-presented pairings (t_25,1_ = 5.45, p < 0.001, Cohen’s d = 1.07). No activation of the uncued S-R pairing was found (t_25,1_ = 1.32, p = 0.2, Cohen’s d = 0.26). As predicted, no differences between cued, uncued and baseline pairings were found in declarative signals (all *t*s < 1.53, all *p*s > 0.14).

**Figure 5.**
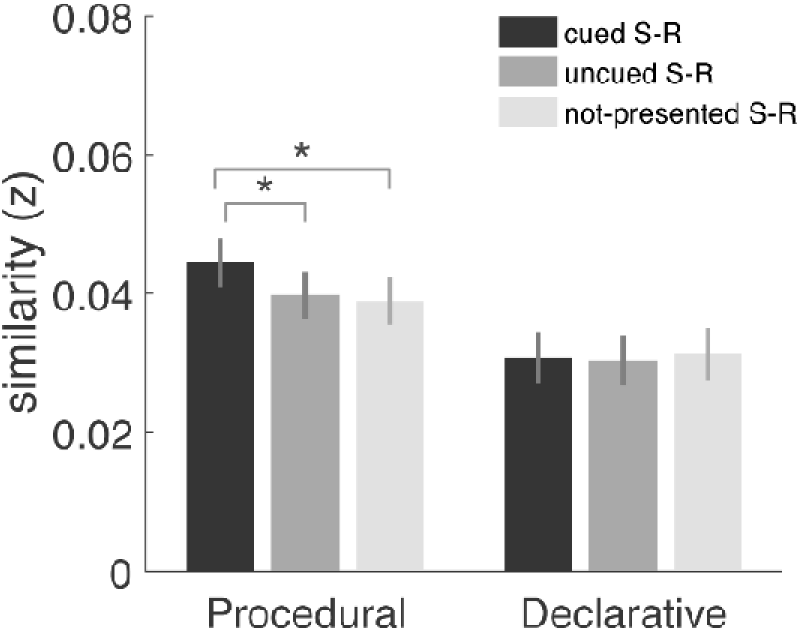
Template tracking procedure results in the primary motor cortex. Bars represent the normalized semi-partial correlation between task data and the procedural and declarative templates of cued, uncued and not presented S-R pairings. Error bars denote within-participants s.e.m (Morey, 2008). Asterisks denote significant differences (p < 0.05, paired t-test).

To further assess the sensitivity of our tracking approach, we repeated the analysis on the beta estimates obtained during the encoding screen of the trial, where no differences should be found between cued and uncued mappings. Given the results during the retro-cue period, here we focused on procedural activation scores. We then entered the template activation scores of the encoding and retro-cue events in a repeated-measures ANOVA with the factors S-R type (cued vs. uncued) and Event (Encoding vs. Retro-cue). Importantly, the activation scores entered were the scores for cued and uncued relative to not-presented mappings, therefore not-presented mappings were not included as a separate level in the S-R type factor of the ANOVA. This analysis yielded a significant S-R type * Event interaction (F_25,1_ = 10.61, p = 0.003, η^2^_p_ = 0.3). The interaction effect revealed a difference in activation of cued and uncued S-Rs only during the retro-cue screen (F = 16.68, p < 0.001), whereas no significant differences were found during the encoding screen (F < 1, p = 0.67). Furthermore, it revealed a boost in the activation of cued mappings during the retro-cue, compared to the encoding screen (F = 4.9, p = 0.036). No difference was found for the uncued S-Rs (F = 2.5, p = 0.125), although activation was numerically weaker during the retro-cue (M = 0.001, SD = 0.004) than during the encoding screen (M = 0.003, SD = 0.006). To directly test whether activation for cued and uncued during the retro-cue was greater that during the encoding screen, we performed a new ANOVA in which we introduced direct scores (not relative to baseline), therefore including not-presented S-R as another level of the S-R type factor. This ANOVA confirmed the Event * S-R type interaction (F = 6.71, p = 0.003). Moreover, post-hoc tests (Bonferroni-corrected) revealed, first, that during the retro-cue screen cued S-Rs had higher activation than uncued (t = 4.66, p < 0.001) and not-presented S-Rs (t = 5.58, p < 0.001), whereas uncued and not-presented S-Rs did not differ (t < 1). In contrast, no differences were found between cued, uncued and not-presented S-Rs during the encoding epoch (all ps > 0.11).

### Declarative and procedural contributions to instruction implementation in frontoparietal regions (and beyond)

To elucidate which signals govern implementation in control-related regions, we carried out the template tracking procedure on each FPN region separately. Furthermore, we decided to include the ventral visual cortex (VVC) in this analysis to explore the effect of implementation in higher-order visual regions, since these have been consistently shown to be involved in instruction processing (González-García et al., 2017a; Muhle-Karbe et al., 2017; Palenciano et al., 2019b, 2019a) and our univariate results also revealed their engagement in the current task.

Importantly, our main goal was to assess whether FPN contained procedural and/or declarative signals during implementation and not to compare the strength of these to each other. Therefore, the raw semi-partial correlation of cued pairings in procedural and declarative formats, which could be biased by for instance higher resemblance between the procedural localizer and the main task, was not informative for our purpose (and did not differ between procedural [M = 0.03, SD = 0.014] and declarative signals [M = 0.03, SD = 0.015], t < 1, p = 0.34). Instead, we focused on the comparison of these activations to the within-localizer empirical baselines provided by the irrelevant mappings on each format. Supporting previous results and theoretical models (Brass et al., 2017; Muhle-Karbe et al., 2017), this analysis (Fig. 6a) revealed prioritization involves the representation of relevant information in an action-oriented format in the FPN (two-tail paired t-test against empirical baseline [not-presented pairings], all *t*s > 2.16, all *p*s < 0.04, all Cohen’s *d* > 0.42). Critically, procedural information of cued pairings was significantly more activated than uncued pairings (all *t*s > 2.26, all *p*s < 0.04, all Cohen’s *d* > 0.44). Regarding declarative information (Fig. 6b), parietal nodes of the FPN showed a specific enhancement of declarative information of the cued S-R pairing, compared to the irrelevant (*t*s > 3, all *p*s < 0.005, all Cohen’s *d* > 0.6) and uncued ones (*t*s > 2.16, all *p*s < 0.02, all Cohen’s *d* > 0.49). In contrast, no significant differences were found in frontal nodes between cued and uncued S-Rs, and cued and irrelevant S-Rs (*t*s < 2.06, all *p*s < 0.05, all Cohen’s *d* < 0.4). To further assess this difference between frontal and parietal nodes we performed an ANOVA on the activation scores with the factors ROI (left frontal, right frontal, left parietal, right parietal) and S-R (cued, irrelevant). This yielded a significant ROI*S-R interaction (F_75,3_ = 4.33, p = 0.007, η^2^_p_ = 0.15), revealing that the declarative activation of cued S-Rs was significantly above baseline in parietal (Fs > 9.5, ps < 0.005) but not frontal nodes (Fs < 0.6, ps > 0.28) of the FPN. Another ANOVA but with cued and uncued as levels of the S-R factor revealed a similar profile difference, although the interaction in this case was not significant (F = 2, p = 0.13). Last, an ANOVA with uncued and irrelevant as S-R levels revealed no significant differences in activation between these two levels (F = 1.22, p = 0.28), and this was not modulated by the ROI (F < 1, p = 0.78), suggesting that declarative information of uncued S-Rs was not activated above baseline in any of the FPN nodes.

**Figure 6.**
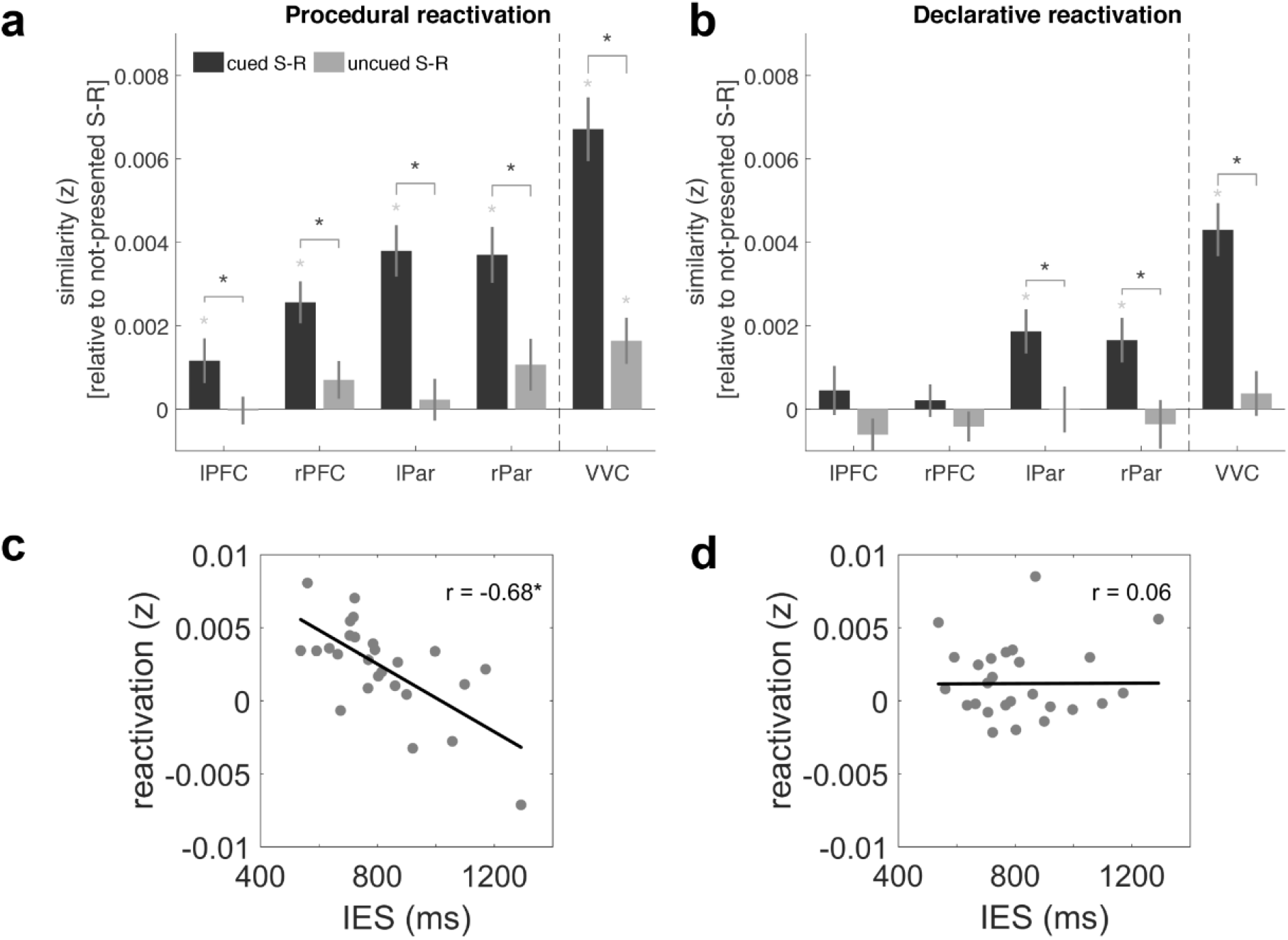
Canonical template tracking procedure results in frontoparietal cortices and ventral visual cortex. Bars represent the normalized semi-partial correlation between task data and (**a**) the procedural and (**b**) declarative templates of cued and uncued S-R pairings. Importantly, raw semi-partial correlation magnitudes of cued pairings are not informative (and did not differ between procedural and declarative signals, all *t*s < 1), and therefore results are plotted relative to the empirical baseline (not-presented S-Rs). Thus, the heights of the bars in panels **a** and **b** simply reflect the difference from baseline and not necessarily different raw semi-partial correlations. Error bars denote within-participants s.e.m. Gray asterisks denote a significant increase from baseline (p < 0.05, paired t-test, FDR-corrected). Black asterisks denote significant differences between cued and uncued pairings (p < 0.05, paired t-test, FDR-corrected). (**c**) Across-participant correlation of Inverse Efficiency Scores and procedural activation index in frontoparietal cortices. (**d**) Correlation of Inverse Efficiency Scores with declarative activation index in frontoparietal cortices. In **c** and **d**, dots represent individual participants, thick lines depict the linear regression fit, and asterisks denote significant Pearson’s correlation (p < 0.05). Activation indices are obtained by subtracting the activation of uncued S-Rs to the activation of cued S-Rs (this can lead to negative values, as can be seen in the **c** and **d**).

Importantly, the lack of declarative activation of cued S-Rs on frontal nodes, and overall the lower enhancement from baseline compared to procedural information (as can be seen comparing Figure 6a and 6b) cannot be due to a lower correlation magnitude of declarative signals with the main task (no significant differences with the correlation magnitude of procedural signals, t < 1, p = 0.45). Still, given the overall low signal-to-noise ratio and pattern reliability in prefrontal cortices (Bhandari et al., 2018), slight differences inherent in the templates could affect the activation measures. For instance, it could be argued that the amount of signal in declarative templates is intrinsically lower than that of procedural templates, which in turn might induce a lack of power to detect the activation of declarative templates in the same regions during the task. To rule out these concerns, for each template and region of the FPN, we compared the signal-to-noise ratio (computed as mean t-value across voxels of the ROI divided by the standard deviation), informational content (computed as Shannon entropy) and correlationability of the templates (i.e. the degree to which individual templates correlated with other templates from the same localizer). This analysis revealed that procedural and declarative FPN templates did not differ in any of these measures (all BF_10_ < 0.5). Moreover, we tested pattern reliability on each localizer separately by assessing the stability of patterns of the same S-R pairing in odd and even trials. To do so, we computed a new GLM with two regressors per S-R pairing, one for odd and another for even trials. We then estimated the correlation (Spearman’s rho) between each regressor. Finally, we compared the similarity of each specific S-R pairing (e.g. in odd trials) with its counterpart (in even trials) to the similarity of the same S-R pairing and the rest of pairings (in even trials). A higher within-pairing compared to between-pairing correlations would suggest reliability of the patterns of activity obtained during the localizers. This analysis revealed statistically reliable patterns in all ROIs and in both localizers (all t > 2.6, all p < 0.05, FDR-corrected for multiple comparisons), supporting the idea that templates contained S-R specific information.

Last, higher-order visual regions showed a similar pattern to parietal nodes of the FPN. As before, the raw semi-partial correlation magnitude of cued pairings with the main task was of no interest and did not differ (t < 1, p = 0.63) between declarative (M = 0.018, SD = 0.024) and procedural signals (M = 0.022, SD = 0.023). Compared to the empirical baseline, we found a significant enhancement of both procedural (*t* = 8.80, *p* < 0.001, Cohen’s *d* = 1.73) and declarative (*t* = 6.76, *p* < 0.001, Cohen’s *d* = 1.33) information of the cued S-R pairing. Crucially, these signals were significantly stronger than the ones of uncued mappings (procedural: [*t* = 6.19, *p* < 0.001, Cohen’s *d* = 1.21]; declarative: [*t* = 5.84, *p* < 0.001, Cohen’s *d* = 1.15]).

### Action-oriented codes support novel instruction implementation

To assess the behavioral relevance of declarative and procedural signals, we reasoned that if action-oriented representations are crucial during implementation in control-related regions, and implementation can be conceived as a behavior-optimized state, then the degree of action-oriented activation should predict the efficiency of instruction execution. To test this hypothesis, we first converted RTs and error rates of informative retro-cue trials into a single compound measure (Inverse Efficiency Scores; IES. IES were obtained by dividing each participant’s mean RT by the percentage of accurate responses (Townsend and Ashby, 1983)). Then, we derived a template activation index by subtracting the degree of activation of cued pairings to that of uncued pairings for each region and format (procedural and declarative). Note that this can lead to a negative activation index (if activation for uncued pairings is stronger than for cued ones). Finally, we correlated individual IES with the activation indices on each region of the FPN. This analysis revealed significant negative correlations in all FPN regions between IES and procedural activation (all Pearson’s *r*s > −0.475, all *p*s < 0.02; See Table 2 for individual ROI Pearson’s rs, p-value and BF_10_ estimates).

**Table 2.** Individual ROI Pearson’s rs, p-values, and BF_10_ estimates. BF interpretation is identical to Table 1.

| | ROI | $r$ | $p$ | $BF_{10}$ |
| --- | --- | --- | --- | --- |
| **procedural** | ldlpfc | -0.475 | 0.014 | 4.203 |
|  | rdlpfc | -0.583 | 0.002 | 24.887 |
|  | lpar | -0.641 | < 0.001 | 88.146 |
|  | rpar | -0.605 | 0.001 | 39.057 |
| **declarative** | ldlpfc | 0.096 | 0.639 | 0.27 |
|  | rdlpfc | -0.339 | 0.09 | 0.955 |
|  | lpar | 0.213 | 0.297 | 0.408 |
|  | rpar | 0.113 | 0.582 | 0.281 |

Regarding declarative codes, we considered three hypotheses. First, if procedural representations are highly dependent on the quality of declarative representations so that participants with high procedural activation also have high declarative activation, one could expect that declarative signals of relevant S-Rs should in principle aid performance as well. Second, declarative activations could be driven primarily by participants with lower procedural activation. In that case, we should find the opposite correlation with behavior (higher declarative activation would predict worse performance). Last, if declarative correlations reflect a residual activation of this coding format that might support the emergence of procedural codes but it is not itself related to behavior, we should expect no correlation. This analysis revealed that IES did not correlate with declarative activation in any region (all *r*s < −0.34, all *p*s > 0.09), although conclusive evidence for the null hypothesis was only found for the left DLPFC and right parietal ROIs (BF10s < 0.3; for the remaining ROIs, evidence was inconclusive; see Table 2).

When averaging activation indices across FPN regions, an identical pattern was found, namely, a significant correlation of IES with procedural (*r* = −0.679, p < 0.001) but not declarative (*r* = 0.06, *p* = 0.77) activation (Fig. 6c-d). Moreover, these two correlations were significantly different (z = −3.13, p = 0.0018). Similar results were obtained when using RTs (procedural: r = −0.67, p < 0.001; declarative: r = 0.076, p = .71) and error rates (procedural: r = −0.54, p = 0.004; declarative: r = −0.019, p = 0.93) as behavioral measures. Also, when removing participants with negative procedural activation scores (which could reflect the use of suboptimal strategies to solve the task, or noise in the estimation of the neural measures) from the analysis, the correlation with IES remained significant (r = −0.54, p = 0.009), whereas the correlation of declarative activation and IES was not significant (r = −0.17, p = 0.43). Finally, we tested if the degree of procedural activation predicted the degree of declarative activation. This correlation was also not significant (r = −0.17, p = 0.40), and if anything pointed in the direction that participants with higher procedural activation were the ones with weakest declarative signals, and vice versa.

Altogether, these results show that the more implementation was governed by relevant procedural codes in the FPN, the faster and more accurately participants executed the instruction. In contrast, the strength of declarative signals of the same S-R association did not predict behavioral performance.

## DISCUSSION

In the current study, we report a pervasive effect of novel instruction implementation across behavioral and neural data. A canonical template tracking procedure revealed that unique declarative and procedural representations govern FPN activity during implementation, prior to execution. These representations were specific to prioritized S-Rs and did not take place for irrelevant mappings. Critically, our results show that procedural (but not declarative) activation in the FPN predicted efficient execution of novel instructions.

### Frontoparietal flexible coding of novel S-Rs

Previous research has highlighted the important role of the FPN in the implementation of novel instructions (Bourguignon et al., 2018; Demanet et al., 2016; González-García et al., 2017a; Hartstra et al., 2011; Muhle-Karbe et al., 2017; Palenciano et al., 2019a, 2019b; Ruge and Wolfensteller, 2010). Accordingly, our results show that the FPN represents relevant S-R pairings during implementation. However, these results remain agnostic regarding the functional nature of the neural codes underlying this effect. Here, we leveraged a canonical template tracking approach to approximate to process-pure measures of procedural and declarative coding formats. This allowed us to later investigate the unique contribution of each format to instruction implementation.

In accordance with the serial-coding hypothesis, we observed that implementation engaged the activation of procedural representations (Brass et al., 2017; Muhle-Karbe et al., 2017). Interestingly, our results show that, in addition to procedural codes, some nodes of the FPN preserve relevant declarative information about the upcoming task.

A first consideration concerns the exact nature of the reactivated signals. In the declarative localizer, participants had to remember specific S-R associations and match them to another S-R probe. In contrast, in the procedural localizer, participants’ goal was to execute the correct response associated with a target stimulus. The different readout from WM thus encouraged different strategies, as suggested by previous studies (González-García et al., 2020; Liefooghe et al., 2012; Muhle-Karbe et al., 2017). Therefore, it is conceivable that templates will contain unique information: a persistent maintenance of the memoranda in the declarative localizer, and a proactive action-oriented representation in the procedural localizer. However, procedural and declarative representations likely share further information, for instance, related to specific perceptual stimulation and domain-general processes, such as arousal or attention. We took several measures to reduce the influence of such components. First, template activation was derived from semi-partial correlations between data from the main task and the localizers. Thus, our measure reflects unique shared variance between the task and the representation of an S-R pairing in a given localizer, partialling out the variance explained by the representation of the same S-R in the remaining localizer. Importantly, our study was aimed at assessing the presence (or lack thereof) of procedural and/or declarative signals and not at comparing to what extent one signal might be more predictive than the other, and therefore we base our results in activation of templates relative to empirical baselines provided by not-presented S-Rs. Second, templates were built for S-R pairings rather than unique mappings, and therefore a contribution of perceptual features to template activation seems unlikely. Moreover, semi-partial correlations were computed between data from the retro-cue screen (in the main task), and inter-stimulus interval (in the localizers), which reduces the likelihood of significant correlations due to perceptual similarity between templates and specific S-Rs. Therefore, although other non-mutually exclusive explanations cannot be fully discarded (e.g. “procedural” templates containing procedural signals but also any other code present in the procedural localizer and not in the declarative one), we believe it is the most parsimonious interpretation to consider that our procedure succeeded at tracking format-specific signals, especially given the validation results in the motor cortex.

An important aspect then concerns the specific functional significance of each format. Regarding procedural templates, although the configuration of the procedural localizer was similar to the main task, the highly action-oriented encoding format encouraged during this localizer was strategically optimal only after the selection process elicited by the retro-cue in the main task. Thus, this localizer allowed us to test whether the selection of an S-R from WM engaged the same procedural signals elicited by encoding tasks with the intention to implement. With respect to the declarative templates, an intriguing question is what exactly is being reactivated, and how is this not present in the procedural localizer (which necessarily has to contain some declarative information as well (Formica et al., 2020a)). One possibility is that the specific demands of each localizer encourage differentiated coding strategies, that is, different readouts from WM could modulate the specific way in which each format is represented. However, we believe a more likely, non-mutually exclusive possibility regards the previously mentioned distinction between the procedural localizer and the main task. Given that the process of maintenance prior to selection is likely diminished in the procedural localizer, it is feasible that such maintenance signals are present in the main task relatively independent from the codes established in the procedural localizer. In turn, it is possible that declarative codes account at least partially for such maintenance components, leading to the observed declarative activations in the main task.

From this standpoint, our results suggest that during novel instruction implementation, FPN regions contain information about the declarative memoranda conveyed by the instruction and an independent action-oriented S-R code that primarily drives task execution.

### Heterogeneous S-R coding within the FPN

Although we did not have specific hypotheses for the role of individual FPN regions, a second important finding concerns the heterogeneity of results within this network. Frontal nodes showed the implementation profile predicted by the serial-coding hypothesis, namely, a primarily procedural representation of instructed content. This is in line with previous studies that propose a crucial role of the frontolateral cortex in the integration of stimulus and response information into a task set based on verbal instructions (De Baene et al., 2012; Hartstra et al., 2012, 2011), as well as in representing task rules (Jackson and Woolgar, 2018; Loose et al., 2017; Wisniewski et al., 2019; Woolgar et al., 2015) and goals (Muhle-Karbe et al., 2014).

In contrast, parietal nodes carried both procedural and declarative information in their patterns of activity. Whereas the role of parietal regions in representing goals and task set information is widely acknowledged (González-García et al., 2017a; Jackson and Woolgar, 2018; Muhle-Karbe et al., 2017, 2014; Palenciano et al., 2019b; Wisniewski et al., 2015; Woolgar et al., 2015), it is unclear what drives such declarative activation. One possibility is that it reflects a category-specific top-down selection scheme, driven by increased attention towards the cued S-R (Nobre et al., 2004; Tamber-Rosenau et al., 2011). The fact that a similar pattern was found in higher-order visual regions, which usually coordinate with parietal cortices to represent relevant task dimensions in anticipation of future demands (González-García et al., 2015; Kuo et al., 2014; Lepsien and Nobre, 2007), further supports this possibility. This tentative interpretation would be coherent with goal neglect effects reported in patients with frontal lobe damage (Duncan et al., 1996). These patients are capable of selecting, maintaining, and remembering task-relevant information, yet their ability to transform relevant information into goal-driven actions is impaired. Such dissociation goes at least partially in line with our results in that (1) goal-oriented representations depends critically on prefrontal cortices (impaired in goal neglect patients), and (2) the involvement of other control-related regions, intact in these patients, boosts the declarative representation of specific task information, such as particular S-R pairings, presumably in coordination with posterior category-selective regions. However, these results should be interpreted with caution, since the difference between frontal and parietal regions could partly reflect a difference in activation magnitude not captured by our method, due to a generally weaker coding in frontal lobes (Bhandari et al., 2018). Still, the impact of this alternative interpretation seems relatively limited, given we observed a similar raw semi-partial correlation magnitude of cued pairings with the main task, and no differences in terms of signal-to-noise ratio, informational content, and correlationability of the templates.

### Implementation as a selective output gating process

Remarkably, although we found both signals in the FPN during implementation, only procedural representations predicted efficient behavior and, if anything, stronger procedural activations did not predict stronger declarative signals. The fact that implementation is signaled by retro-cues renders this effect relevant to current debates on information prioritization and WM architecture. In this regard, our results are consistent with the interpretation of implementation as a particular instance of output gating mechanisms. Similar to the idea of an input gate that limits what information enters WM, some computational models propose an additional gate that determines which pieces of this information will drive behavior (Chatham et al., 2014). Recent theoretical frameworks suggest a role of prioritization not only in selecting relevant content from WM but also in reformatting such content into a “behavior-guiding representational state” (Myers et al., 2017), analogous to an output gating mechanism. Interestingly, these models propose that whereas other control-related regions might be involved in attention-driven representations of relevant content, frontal regions are thought to be especially important in transferring this content into a state that is optimal for behavior. Accordingly, our results suggest that an action-oriented representation of novel instructions dominates activity in frontal cortices and that this representational format is tightly linked to behavioral efficiency. A limitation of the current study concerns the lack of specificity on what precise information is captured on each template: it is possible that part of the correlation with behavior we observe is driven not only by procedural codes but also by any other code of different nature that is present in the procedural localizer and not in the declarative one, although what this code would be specifically remains unknown. This question awaits further investigation.

Importantly, our results reveal that the neural substrate of instruction prioritization involves further brain regions, such as category-selective and parietal cortices, and that procedural and declarative information coexist in these regions. This raises the question of what the contribution of declarative representations might be. One possibility is that declarative codes support the generation and maintenance of procedural codes, but once these are created, they do not directly contribute to behavior. It should be noted, however, that fMRI data lacks the temporal resolution to discern the dynamic profile of these two representational formats. Thus, the conclusions about the dynamics of declarative and procedural codes in the FPN we can extract from the current dataset are limited. Further research is needed to elucidate whether, in smaller timescales, a temporal hierarchy between these two signals can be established or, in contrast, whether both signals are held simultaneously in these regions. Future studies should employ time-resolved techniques that can succeed at characterizing the contribution of different brain regions to separate control and WM processes (Quentin et al., 2019).

Last, the current work relies on a relatively high number of tests and decisions along the analysis pipeline, which could potentially impact the results and the conclusions extracted from them (Botvinik-Nezer et al., 2020). As such, the new method proposed here would benefit from independent conceptual replications and extension of the current findings in the future.

## CONCLUSIONS

In summary, the present study reveals the strong impact of instruction implementation on frontoparietal regions. We observed that these regions contain information about prioritized S-R pairings in detriment of the irrelevant ones during implementation. This information contained two non-overlapping neural codes, one reflecting the declarative maintenance of task, and another, more pragmatic, action-oriented coding of the instruction. Importantly, the strength of procedural activation predicted behavioral performance. Altogether, our results highlight the contribution of frontoparietal regions to output gating mechanisms that drive flexible behavior.

## Financial interests or conflicts of interest

none declared.

## Acknowledgments

C.G.G. and S.F. were supported by the Special Research Fund of Ghent University BOF.GOA.2017.0002.03. C.G.G. was additionally supported by the European Union’s Horizon 2020 research and innovation programme under the Marie Sklodowska-Curie grant agreement no. 835767. D.W. was supported by FWO and the European Union’s Horizon 2020 Research and Innovation Program under the Marie Skłodowska-Curie grant agreement no. 665501. We thank Senne Braem for feedback on previous drafts of the manuscript.

## Notes

### Competing Interest Statement

The authors have declared no competing interest.

